# Proteomic analyses of the Arabidopsis cap-binding complex define core set and TOR-dependent protein components

**DOI:** 10.1101/2023.02.16.528822

**Authors:** Annemarie Matthes, Laurence Ouibrahim, Cécile Lecampion, Carole Caranta, Marlène Davanture, Régine Lebrun, Yohann Couté, Mathieu Baudet, Christian Meyer, Christophe Robaglia

## Abstract

The eukaryotic cap-binding complex (CBC) is a hub for regulations affecting mRNA behaviour including translation, degradation and storage. Beside the core eukaryotic translation initiation factors, other proteins, many of which are yet unknown, are thought to interact stably or transiently with the CBC depending on cell status. The prototype of these regulators is the animal eIF4E binding protein (4E-BP), a direct target of the TOR (Target of Rapamycin) kinase that competes with the cap-binding protein eIF4E, thus repressing translation. In plants, no functional homologs of 4E-BP have so far been characterized. In this work we performed several deep proteomic analyses of the Arabidopsis CBC after cap-affinity purification from wild-type plants. We also investigated the CBC in eIF4E mutant plants, Arabidopsis lines with lower TOR activity, or during infection with eIF4E-dependent potyviruses, conditions which are all affecting translation at the initiation level. These analyses allowed us to define a limited core set of CBC components, which were detected in all samples. Interestingly, we identified proteins, like AGO1 or VCS, which were always detected in conditions where either TOR or mRNA translation were reduced. Meta-analysis of these data revealed several new plant interactors of the CBC, potentially defining pathways related to mRNA stability and degradation, metabolism and viral life cycle. A search for eIF4E binding motifs identified several new potential 4E-BP relatives in plants.

## Introduction

Protein synthesis is one of the cell’s most energy consuming processes and is thus regulated at many levels and particularly at the level of the initiation of mRNA translation (Shah et al, 2013). Most eukaryotic mRNA are terminated at the 5’ end by a 7-methyl guanylate nucleotide (m7GpppN), called the cap, which is recognized by a conserved protein complex, (cap-binding complex, CBC) mediating access to the ribosomal 40S subunit. The CBC is structured by the eIF4G scaffold that bridges the eIF4E cap-binding protein with the eIF4A helicases, the multi-subunit eIF3 and the 40S ribosomal subunit, to form the 43S translation preinitiation complex in association with mRNA (Jackson et al, 2010; Browning and Bailey-Serres, 2015; Kumar et al, 2016). eIF4E and eIF4G are thought to be the ancestral forms, while in flowering plants a distinct CBC is structured by eIF(iso)4E and eIF(iso)4G isoforms (Patrick and Browning, 2012). The CBC is a critical point for modulation of mRNA fate across diverse eukaryotic species. Several physiologically regulated proteins were found to interfere with the interaction between eIF4E and eIF4G modulating translation efficiency. Among the best-known examples are eIF4E-binding proteins (4E-BPs) found in several metazoan and fungal species (Mamane et al, 2006). 4E-BPs are phosphorylated by the highly conserved Target of Rapamycin (TOR) kinase, itself activated by growth promoting factors such as sugars, amino-acids and hormones (Condon and Sabatini, 2019; Ingargiola et al, 2020). In its phosphorylated state, 4E-BP does not interfere with eIF4E to eIF4G binding, thus allowing protein synthesis to proceed freely, promoting cell proliferation and growth. In the dephosphorylated state, when TOR is inactivated by nutrient starvation or stress conditions, 4E-BP sequesters eIF4E to prevent 43S complex formation, leading to global repression of translation and the promotion of cell survival and quiescence. The existence of regulators similar to animal 4E-BPs was for a long time debated in plants. Recently, two Arabidopsis eIF4E protein interactors CBE1 and CERES were identified but their functions seem different from the ones of animal 4E-BPs. CBE1 was found to be required for proper cell cycle gene expression (Patrick et al, 2018), and CERES was proposed to replace eIF4G allowing the formation of an alternative 43S preinitiation complex (Toribio et al, 2019). Moreover, the link between these proteins and the TOR pathway has not been clearly established, although the phosphorylation of CBE1 was found to be regulated by TOR activity (Scarpin et al, 2020). An Arabidopsis ortholog of the GIGYF alternative CBC component was also found to be a target of TOR (van Leene et al, 2019). It is known that mammalian LARP1 protein (La-related protein1) binds cap analog and blocks eIF4F formation (Lahr et al, 2017). The Arabidopsis orthologs of LARP1 have been shown to be involved in the regulation of mRNA stability after heat stress (Merret et al, 2013) and to be phosphorylated in a TOR-dependent manner (van Leene et al, 2019; Scarpin et al, 2020).

The critical role of the CBC in translational regulations is further highlighted by its frequent targeting by viruses. The Hantavirus N protein replaces eIF4E and eIF4G attracting the ribosome to the viral RNA (Mir and Panganiban, 2008). Picornaviruses infection leads to eIF4G cleavage promoting shut-off of host mRNA translation, while translation of the viral RNA is favored by the presence of an Internal Ribosome Entry Sites (IRES). The Potyviral 5’ genome linked protein (VPg) binds eIF4E and eIF(iso)4E promoting viral translation (Khan et al, 2008; Eskelin et al, 2011). A specific structure at the 3’ end of several plant RNA viral genomes (3’-cap-independent translation enhancer or 3’CITE) can also directly recruit eIF4E and stimulate viral translation (Simon and Miller, 2013). eIF4E proteins have repeatedly evaded their capture by viral proteins or RNA, providing viral resistance in several plant species (Robaglia and Caranta, 2006; Nieto et al, 2006; Poulicard et al, 2016)

Purification and biochemical analysis of the CBC can be performed through affinity purification to resin-coupled m7GTP, which acts as a cap analogue (Sonenberg et al, 1979). Copurifying proteins were found to be considerably enriched for proteins involved in translation initiation and in the regulation of protein synthesis in animals as well as in plants. Indeed, in an initial study analyzing the composition of the CBC in Arabidopsis, about 20 new proteins not previously known to be part of the Arabidopsis CBC were identified (Bush et al, 2009). Several of these candidates were later confirmed as genuine components of the CBC which could regulate translation such as the CBE1 protein (At4g01290, Patrick et al, 2018), the GRP8 RNA binding glycine rich protein (At4g39260) involved in RNA splicing and flowering (Steffen et al, 2019) and the EXA1 GIGYF-like protein (At5g42950), involved in RNA virus replication and defense gene expression (Hashimoto et al, 2016; Wu et al, 2019).

In the present work we took advantage of the recent progresses in the sensitivity, speed and resolution of mass spectrometers to perform a wide analysis of affinity-purified CBC proteomic data from Arabidopsis plants. One of the main goals of this study was to identify the most stable and ubiquitous components of the Arabidopsis core CBC by identifying proteins which were always present in the proteomic analyses. We present a detailed analysis of the CBC in wild-type (WT) Arabidopsis and compared it to several lines where translation initiation is affected by different causes. These are a mutant devoid of the eIF4E protein, plants infected by the Watermelon Mosaic potyvirus (WMV) and plants where the TOR pathway was repressed by genetic or chemical repression. This wide study allows us to identify invariant components of the CBC in plants as well as those that appear in the CBC only upon TOR repression or viral infection. These proteins could provide new leads for the study of translational regulation in plants.

## Materials and Methods

### Plant material and growth conditions

Experiments are summarized in Table I. Arabidopsis Col0 ecotype was the WT background for most experiments excepted for experiments with TOR inducible RNAi lines which were performed with Landsberg *erecta* ecotype. TOR RNAi constitutive line 35.7, TOR RNAi inducible lines 5.2 and 6.3 and GUS control line were described in Deprost et al, 2007. The eIF4E KO mutant line SALK-145583 was described in Bastet et al, 2018.

**Table I:** Details of experimental procedures for proteomic analyses of the Arabidopsis cap-binding complex. The total number of proteins detected in each experiment in indicated. Experimental data can be found in: G1 to G4: supplemental data tables 2-5 G5 to G6: supplemental data tables 6-7 M1-3: supplemental data table 8 L3 to L4: supplemental data table 9-10

| Name of experiment | Genotype/inhibitor | Growth conditions | Purification method | Total number of detected proteins | Platform |
| --- | --- | --- | --- | --- | --- |
| G1: Control eIF4E mutant TOR RNAi | Col0 | 8d seedlings liquid culture | m7GTP sepharose, gel separation. The gel was cut in two slices which were analyzed independently. | 486 | Edyp-Grenoble |
| G2: eIF4E mutant | Col0 eIF4E-/- | 8d seedlings liquid culture | m7GTP sepharose, gel separation. The gel was cut in two slices which were analyzed independently. | 345 | Edyp-Grenoble |
| G3: Constitutive TOR RNAi | Col0 TOR constitutive RNAi | 8d seedlings liquid culture | m7GTP sepharose, gel separation. The gel was cut in two slices which were analyzed independently. | 190 | Edyp-Grenoble |
| G4: Control unspecific binding | Col0 | 8d seedlings liquid culture | Sepharose4B, gel separation. The gel was cut in two slices which were analyzed independently. | 354 | Edyp-Grenoble |
| G5 : Control WMV infection | Col0 | 35d old plants, soil | m7GTP sepharose | 165 | Edyp-Grenoble |
| G6: WMV infection | Col0+WMV | 35d old plants, soil | m7GTP sepharose | 111 | Edyp-Grenoble |
| M1: Control TOR inducible RNAi | Le alca::GUS | 8d seedlings liquid culture | m7GTP sepharose | 86 | PAPPSO |
| M2: TOR inducible RNAi 5.2 line | Le alca::TOR-IRNAi 1 | 8d seedlings liquid culture | m7GTP sepharose | 181 | PAPPSO |
| M3: TOR inducible RNAi 6.3 line | Le alca::TOR-IRNAi 2 | 8d seedlings liquid culture | m7GTP sepharose | 183 | PAPPSO |
| L3: AZD8055 treatment | Col0 | 21d seedlings hydroponic | Sepharose4B preadsorption, m7GTP sepharose. The gel was cut in ten slices which were analyzed independently. | 822 | Marseille IMM |
| L4: Control DMSO treatment for AZD8055 treatment | Col0 + AZD8055 | 21d seedlings hydroponic | Sepharose4B preadsorption, m7GTP sepharose. The gel was cut in ten slices which were analyzed independently. | 997 | Marseille IMM |

For experiments G1 to G4, and M1 to M3, 100mg sterilized seeds were grown in liquid modified Hoagland medium, with agitation for 8 days after germination before either induction or harvesting. Induction of the TOR RNAi constructs were performed by adding ethanol to 0.1% and overnight incubation before harvesting (Deprost et al, 2007).

Experiments G5 and G6 were performed with plants grown in soil for 35d. Watermelon Mosaic Virus inoculation was performed by carborundum abrasion of four leaves of each plant at 28d with phosphate buffer (Ouibrahim et al, 2015).

For experiments L3 and L4, plants were grown hydroponically in boxes filled with 100ml of modified Hoagland medium. The media was changed every week and AZD-8055, a strong and specific TOR kinase inhibitor (Montané and Menand, 2013), was added directly to the medium at 1 uM in DMSO, after 3 weeks of culture followed by 15 min incubation (sample L3). Controls boxes received and equal volume of DMSO (sample L4).

### Cap-binding affinity purification

Plant material was blotted dry, weighted and frozen in liquid N2, finely grounded before addition of extraction buffer (1mg/ml) and homogenization. Extraction buffer was 40 mM HEPES buffer containing 0.1M KCl, 10% (v/v) glycerol, 1 mM DTT, 0.3% (w/v) CHAPS containing protease inhibitors (1 tablet Biorad/10ml ; 1/100 v/v PMSF). Phosphatase inhibitors were also added in L3 and L4 experiments. Extracts were clarified at 15000 rpm at 4°C in SW41 rotor for 10 min in a Beckman ultracentrifuge, protein concentration in the supernatant (usually around 2mg/ml) was measured using the Bradford assay. Two mg of proteins in buffer were mixed with 50ul 7-Methyl-GTP-Sepharose 4B (GE Healthcare) previously equilibrated with extraction buffer, and incubated overnight at 4°C on a rotating wheel, before being loaded onto mini-columns. The columns were washed with 4ml of extraction buffer, then with 1ml of extraction buffer containing 0.1 mM GTP, elution was done with 50-100ul of 5mM m7GTP. In experiments L3 and L4, the supernatant was preincubated with Sepharose 4B for removing proteins binding unspecifically to this support. Experiment G4 was performed with plain Sepharose 4B instead of 7-Methyl-GTP-Sepharose 4B. Samples were either sent and processed directly by the platforms or proteins were separated on polyacrylamide gels (10%) and lanes were cut out, digested by trypsin and analyzed by MS/MS as previously described (Dobrenel et al. 2016). Briefly, peptides obtained after trypsin digestion were analyzed by nano LC-MS/MS on a NanoLC-Ultra system (Eksigent). Eluted peptides were analyzed with a Q-Exactive mass spectrometer (Thermo Electron) using a nano-electrospray interface (non-coated capillary probe, 10 μ i.d; New Objective). Peptides and the corresponding proteins were identified and grouped with X!TandemPipeline using the X!Tandem Piledriver (2015.04.01) release and the TAIR10 protein library with the phosphorylation of serine, threonine and tyrosine as a potential peptide modification. Precursor mass tolerance was 10 ppm and fragment mass tolerance was 0.02 Th. Identified proteins were filtered and grouped using the X!TandemPipeline v3.3.41. Data filtering was achieved according to a peptide E-value lower than 0.01. The false discovery rate (FDR) was estimated to 0.92%.

Samples were processed and analyzed by core facilities: Edyp http://www.edyp.fr/web/ (G1 to G6); PAPPSO http://pappso.inrae.fr/ (M1 to M3); Marseille proteomique https://marseille-proteomique.univ-amu.fr/ (L3 and L4) (Table 1). During all procedures great care was exerted to minimize contamination by human or animal keratins.

### Accession to the raw data

The raw data have been deposited to the ProteomeXchange Consortium via the PRIDE (Perez-Rivero et al, 2019) partner repository under the following dataset identifiers:

Experiments G1 to G6: PXD028376

Experiments M1 to M3: PXD028533

Experiments L3 and L4: PXD029102

### Polysome preparation and analysis

Polysomes were prepared and fractionated exactly as described by Lecampion et al, 2016.

## Results and discussion

### Core proteins of the Arabidopsis cap-binding complex

Four different sets of proteomic analyses were performed that were named according to the proteomic core facility performing the analysis as well as to the different conditions or mutants being analyzed, i.e. Grenoble facility (G1-6) for analysis of TOR constitutive RNAi, eiF4E mutants and WMV virus infection, Moulon (M1-3) for analysis or TOR RNAi inducible lines and Marseille (L3 and L4) for plants treated with AZD-8055 a TOR inhibitor (see Table I for details). The objective was to define a robust CBC core protein set from various analysis procedures. Different conditions potentially affecting CBC composition were used, each set including a control experiment based either on the Arabidopsis Col-0 ecotype or the Landsberg *erecta* ecotype (M1-M3). In experiments G1-4 a control where proteins were captured on Sepharose-4B without attached cap-analog was performed to evaluate non-specific binding. In experiments L3 and L4, proteins extracts were preincubated with Sepharose-4B. Between 86 to 997 proteins were identified per experiment (Table I, Supplementary Table 1). For the experiments L3 and L4, the polyacrylamide electrophoresis gel was cut in ten slices which were analyzed separately. Specific proteins lists were then assembled to generate the total list of proteins in which duplicates were removed.. The majority of proteins were identified in only one or two proteomic analyses consistent with the idea that the composition of the CBC is dynamic by nature and that it can be influenced by experimental context and growth conditions, but also by the treatment of mass spectrometry data and instrument performance (Supplementary Table 1 to 10). We first performed a comparison of all lists of proteins found in control CBC experiments by defining their intersections to identify common components.

We thus established a core set of nine common proteins identified in all proteomic analyses (Table II, Figure 1). Seven out of these nine proteins were previously identified in a study of the composition of the CBC in Arabidopsis cell lines (Bush et al., 2009). These proteins represent canonical components of the CBC including eIF4E, eIF(iso)4E, eIF(iso)4G and nuclear CBP20 and CBP80 (Castellano and Merchante, 2020). Another conserved protein is the RNS2 RNase (At2g39780) which has been shown to be involved in rRNA recycling and whose mutation results in reduced TOR activity (Kazibwe et al., 2020). A protein containing a GDSL-motif lipase (At1g29670) which interacts with SCE1 and Sumo, both of which interact with eIF3G, according to the Biogrid database (https://thebiogrid.org/) was also systematically identified. Almost all experiments, except the AZD-8055 control analysis, identified the beta subunit homolog potassium channel (At1g04690) which interacts with TAP46, a component of the PP2A phosphatase complex that is a target of the TOR kinase (van Leene et al., 2019; and reviewed in Ingargiola et al, 2020). The canonical eI4FG factor was often detected but less robustly than eIF(iso)4Gs, which could reflect a weaker association with eIF4E. Nitrate reductase isoforms (NIA1 At1g77760; NIA2 At1g37130) were identified in several experiments and interact with the eIF2B-delta initiation factor that was also found to interact with the TORC1 complex (Figure 1; van Leene et al. 2019). The CERES protein (At4g23840), described as an eIF4E interacting protein (Toribio et al., 2019), was found in CBCs purified from hydroponics cultures (L4) but not after AZD-8055 treatment. In Brassicaceae, two recently duplicated genes can potentially code for eIF4EB (At1g29550) and eIF4EC (At1g29590) isoforms of eIF4E (Patrick and Browning, 2012). However, those proteins were not detected in our experiments. This confirms previous observations that eIF4EC appears unexpressed and that eIF4EB is expressed at low level (Patrick et al, 2014). The different subunits of eIF3E (eIF3a, eIF3c, eIF3e, eIF3f, eIF3h) were also frequently identified but at a lower frequency than core components.

**Table II:**
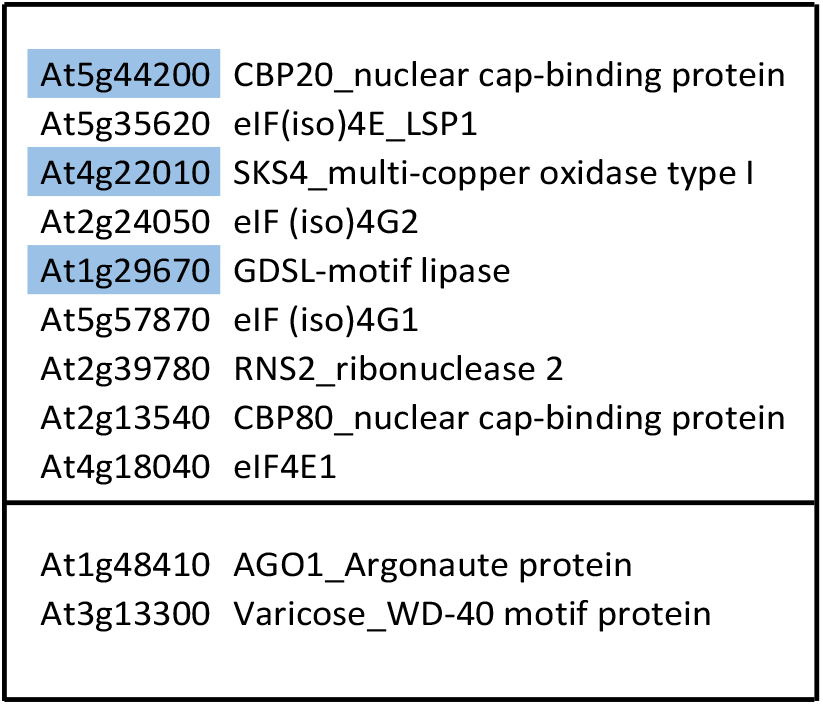
Proteins of the core CBC present in all proteomic analyses. Upper panel: Core CBC defined by proteomic analyses of the control Col-0 line. Proteins in blue were not found in the CBC after TOR silencing. Lower panel: proteins present in experiments after TOR RNA silencing or inhibition with AZD-8055.

**Figure 1:**
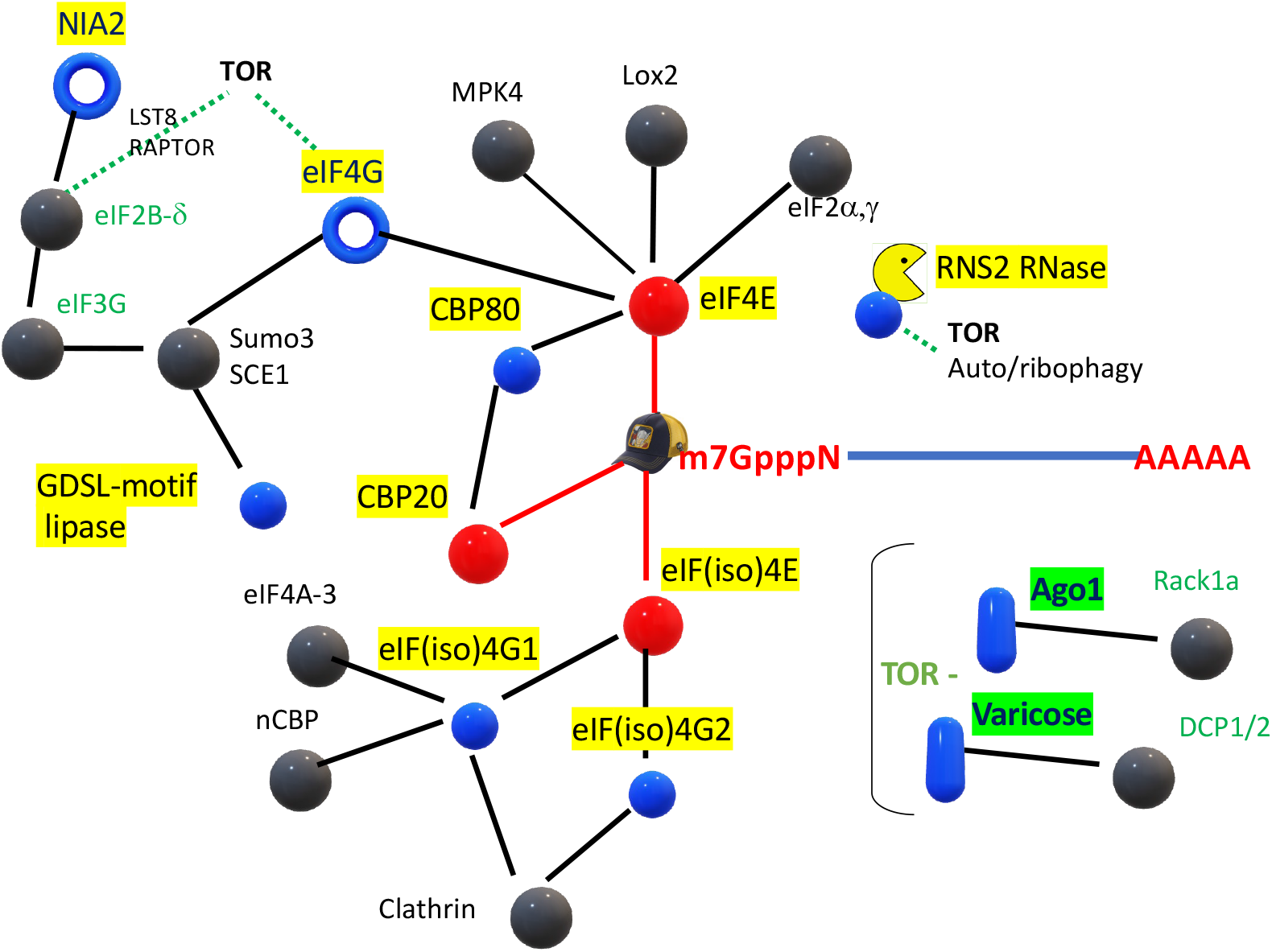
Identification of core CBC and TOR-dependent components. Spheres in blue and red (expected direct interaction with m7GTP, red lines) represent the core CBC found in all control experiments using control Arabidopsis Col-0 plants. Blue donuts show proteins often but not always detected in the proteomic analyses of the CBC. Black spheres and lines represent interacting proteins identified in the Biogrid database Blue cylinders represent the AGO1 and Varicose (VCS) proteins found in all cap-binding affinity purifications when TOR activity was inhibited (RNAi, AZD-8055 treatment).

Several different RNA helicases and RNA-binding proteins were identified in association with the CBC (see Tables II and III). Among them the AGO1 (At1g48410) protein was identified in almost all cases where TOR was inhibited either by silencing or by addition of AZD-8055 as well as in WMV infected plants (4 out of 5 experiments, see discussion below). The AGO2 (At1g31280) protein was also identified in some experiments after TOR inhibition (L3). GRP7 (At2g21660) and GRP8 (At4g39260) proteins are also present in most experiments, they were previously associated with the CBC and their role in RNA metabolism has been established (Steffen et al, 2019).

**Table III:**
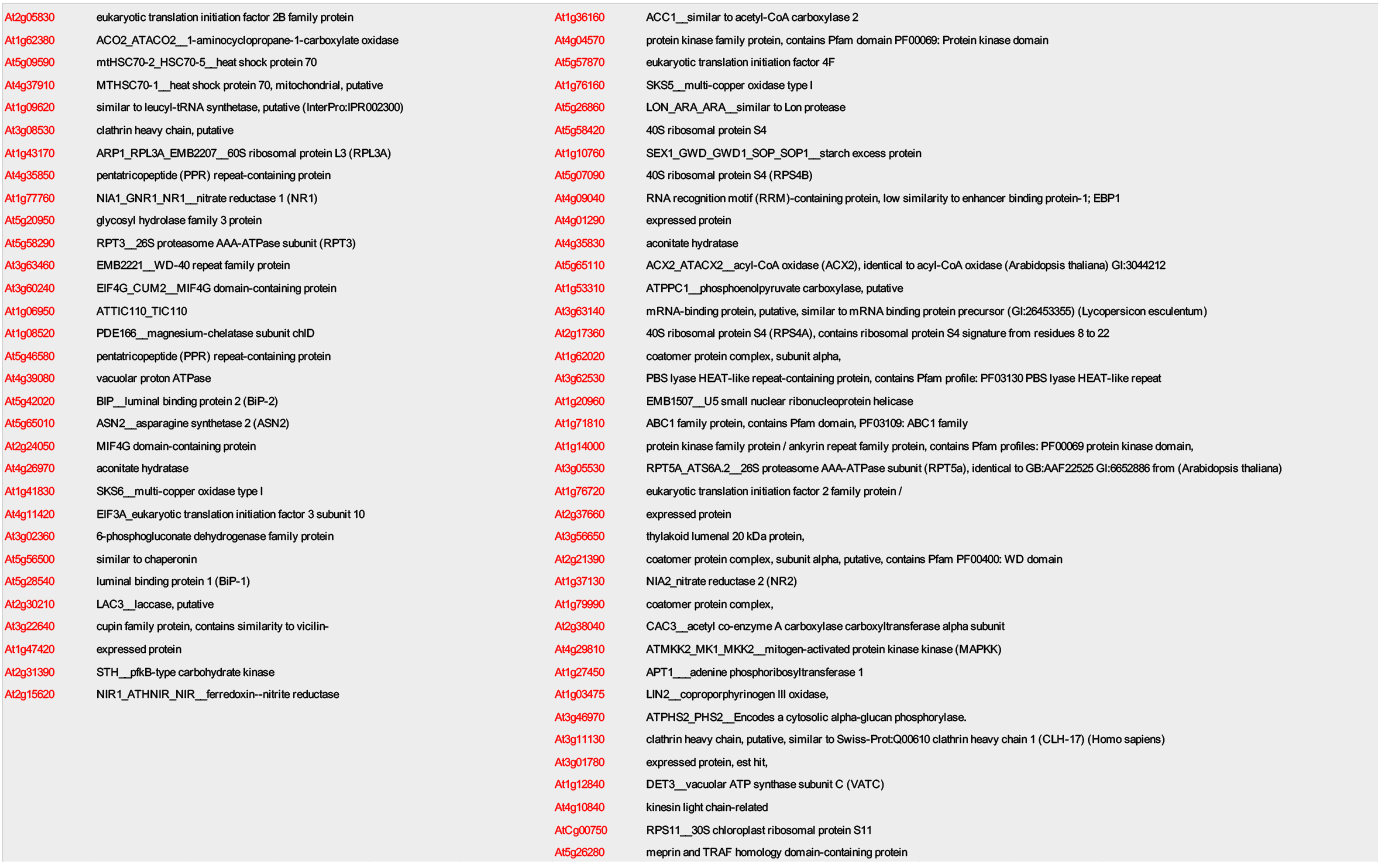
Subset of proteins from all the 11 experiments bearing the modified 4E-BP motif [HRKQE]-x-x-[YRK]-x-[RHST]-x-[FAVLIME]-[LQE]-[MLWFYI]

### The Arabidopsis cap-binding complex in absence of eIF4E

We performed an analysis of the Arabidopsis CBC in the absence of eIF4E. The Salk-145583 mutant line contains a T-DNA inserted in the first intron of the eIF4E gene (At4g18040). The lack of eIF4E in this mutant was previously characterized by western blot analysis (Bastet et al, 2018). Cap dependent translation in this mutant line is thought to rely mostly, if not entirely, on eIF(iso)4E, since *eif4e/eIf(iso)4e* double mutants are not viable (Patrick et al, 2014; Callot and Gallois, 2014) ruling out a substantial contribution of eIF4EB and eIF4EC. The eIF4E KO line exhibits a slow growth phenotype with a delay in flowering (Bastet et al, 2018). Polysome analysis reveal a lower accumulation of polysomes over monosomes compared to WT suggesting a global defect in translation (Figure 2). The absence of eIF4E was further confirmed in the proteomic analysis where it could not be detected in the mutant line (experiment G2), while it was well represented in all other conditions. eIF4E is thought to form a specific complex with eIF4G (At3g60240). Indeed, eIF4G was not detected in the eIF4E mutant line supporting the idea that heterologous eIF(iso)4E/eIF4G complex formation does not occur *in vivo*. In support of this, Patrick and Browning (2014) mention that plants with only eIF(iso)4E and eIF4G are not viable and Patrick et al, (2018) reported that eIF4G was also absent in the CBC of another eIF4E mutant line. Remarkably, AGO1 and NIA1 are still detected in the eIF4E mutant CBC (experiment G2), suggesting that they can associate with the eIF(iso)4E-dependent CBC.

**Figure 2:**
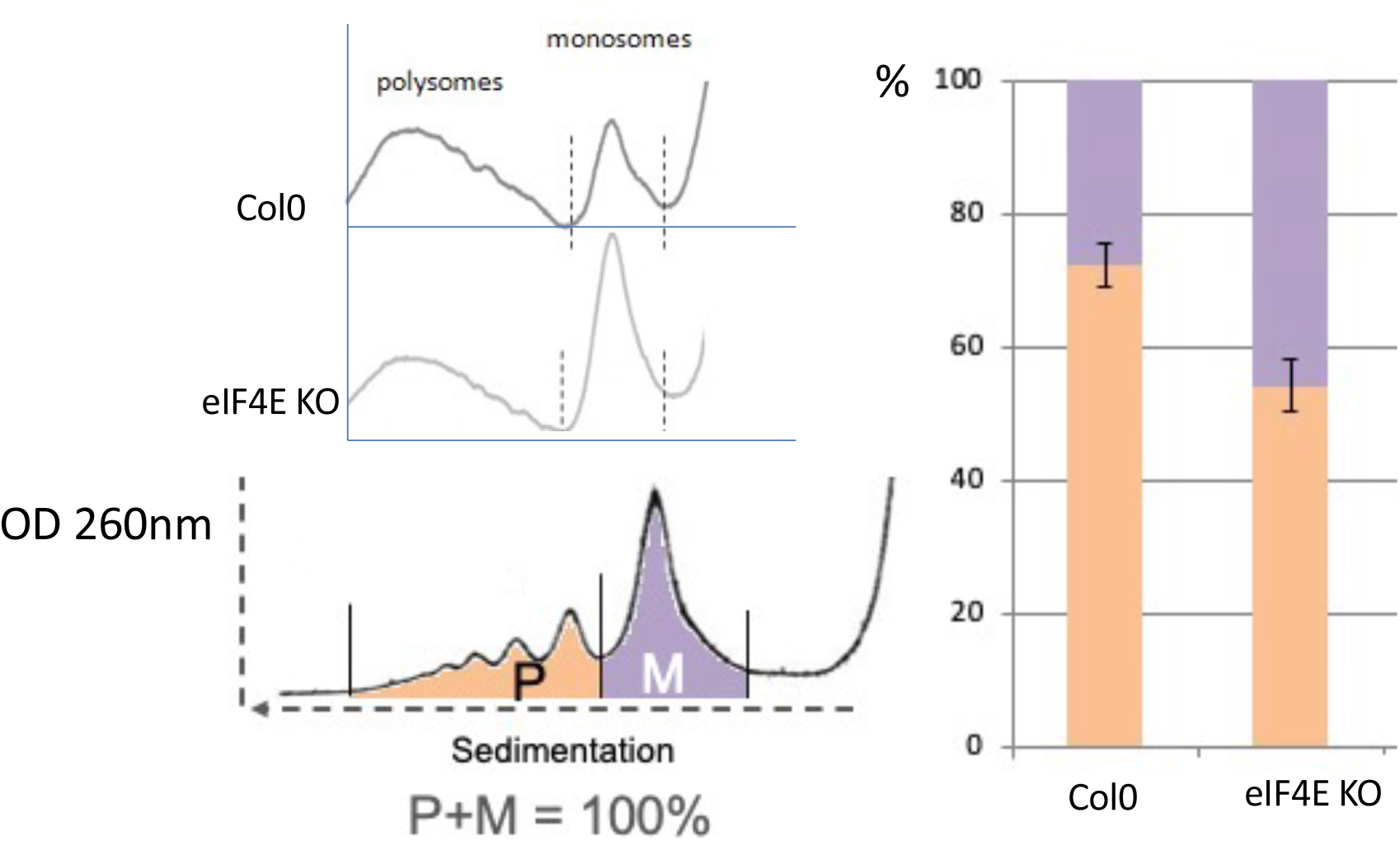
Polysome profiles of Col-0 and eIF4E KO seedlings. The area under the curves of monosomes and polysomes of each profile was integrated to give the graph shown to the right. Orange, polysomes; violet, monosomes.

**Figure 3:**
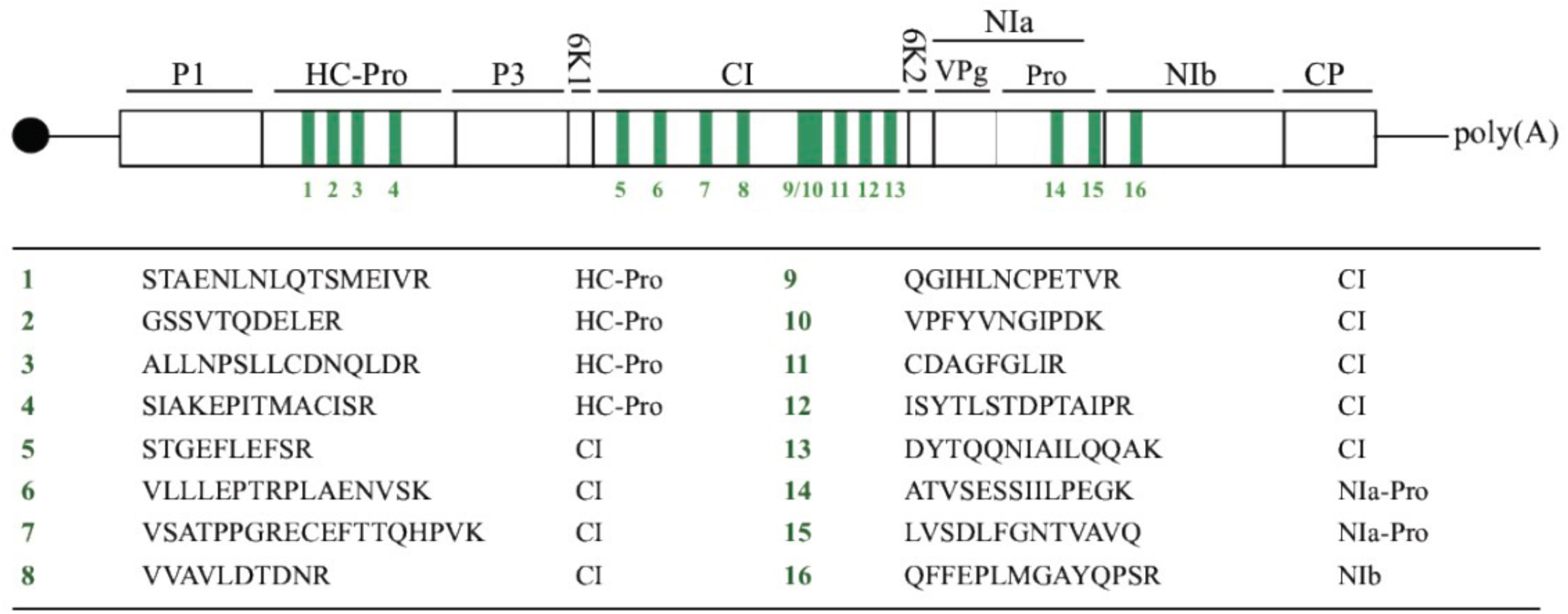
WMV polyprotein specific peptides identified in the CBC. (see Supplemental Data 6 and 7 for details).

### Cap-binding complex modulation during TOR inhibition

Similarly, to the mammalian and yeast TOR pathways, inhibition of the plant TOR repress translation (Deprost et al, 2007). However, the molecular aspects of this repression are not fully understood. For example, functional equivalents of the mammalian 4E-BPs are still elusive in plants. Therefore, CBC analysis during TOR inhibition could reveal mediators of translational repression. To obtain a common picture of the effects of TOR inhibition, we analyzed the CBC in three different conditions: constitutive RNAi (experiment G3) ethanol inducible RNAi (experiments M2 and 3), and treatment with AZD-8055 (experiment L3).

AGO1 is one of the proteins that is consistently detected in the CBC during different conditions of TOR inhibition. AGO1 is responsible of the mRNA cleavage activity of the RNA induced silencing complex (RISC) guided by different classes of small RNA (Song et al, 2019). Independently of RNA cleavage, RNA silencing has as strong effect on mRNA translation that is not completely understood (Brodersen et al, 2008; Lanet et al, 2009; Song et al, 2019). AGO1 has been associated with translating polysomes in plants (Lanet et al, 2009) and in other organisms, such as mammals and insects, miRNA and the RISC complex are known to interfere with the translation initiation step, although the targets and mechanism of action are still controversial (Jonas et al, 2015). The VARICOSE (VCS, At3g13300) WD-40 containing protein, a partner of the plant decapping complex, was also identified in the CBC after TOR inhibition (either by inducible silencing, M2 and M3, or after inhibition by AZD-8055, L3). The decapping complex removes the cap from the 5′ end of mRNAs and is involved in eukaryotic mRNA decay and is composed of Decapping 1 (DCP1), Decapping 2 (DCP2) and VCS. Moreover, VCS can be phosphorylated by SnRK2 kinases (Soma et al. 2017; Kawa et al. 2020) which also inhibit TOR activity by phosphorylating RAPTOR (Wang et al. 2018) and controlling SnRK1 activity (Belda-Palazon et al. 2020). This could indicate a coordinated action of SnRK kinases on both the function of TOR and the stability of mRNAs. VCS was also found to be involved in the regulation of miRNA accumulation, which could suggest that VCS and AGO1 are TOR-dependent interactors of the CBC (Motomura et al. 2012).

### Potential plant 4E-BPs

4E-BPs are involved in TOR mediated translational repression in mammals, but the plants analogs are still unknown. We therefore attempted to use our set of proteins to short-list potential plant 4E-BPs. We started from the known 4E-BP motif (Gosselin et al, 2013) [HRKQ]-x-x-Y-x-[RH]-x-[FAVLIM]-L-[MLWFY] that we slightly modified to better match Arabidopsis 4G and (iso)4G in [HRKQE]-x-x-[YRK]-x-[RHST]-x-[FAVLIME]-[LQE]-[MLWFYI]. A search of the whole Arabidopsis proteome provided 2598 hits. These were then compared to our complete set of proteins co-purified with the CBC providing 69 candidates (Table III). This list includes the CBE1 protein whose absence was previously shown to delay flowering and derepress cell-cycle related genes, a likely TOR output (Patrick et al, 2018). This supports the fact that the short-list contains new potential translational regulators linked to the plant TOR pathway, which would need further studies to establish their roles in regulating eIF4E-dependent translation initiation.

### Potyviral proteins interacts with the CBC

Plant potyviruses form a large family of RNA viruses. Their genome bears a 5’ Viral Protein Genome linked (VPg known to interact with eIF4E and eIF(iso)4E proteins for promotion of viral infection (Robaglia and Caranta, 2006)). Depending on the virus species, eIF4E, eIF(iso)4E or both can be captured by the VPg. To examine whether viral infection affect CBC composition and if other viral proteins besides VPg (or its precursor the Nia protease) could directly or indirectly bind the CBC, we infected young Arabidopsis plants with Watermelon Mosaic Virus (WMV), a potyvirus using both eIF4E and eIF(iso)4E (Bastet et al, 2018) prior to CBC purification and analysis. Besides VPg-pro, we identified the multifunctional protein Hc-Pro, the viral coat protein, the viral polymerase Nib and the Cylindrical inclusion protein (CI). CI, HC-pro and VPg-pro were previously found to form a complex (Roudet-Tavert. et al, 2007, 2012; Zilian et al, 2011). Hc-Pro is a potyvirus specific multifunctional protein with multiple roles in virus/host interaction. One of its cellular functions is to suppress antiviral RNA silencing, although its precise molecular action is not fully understood. As discussed above, RNA silencing is associated with translational repression that likely involve AGO1, the main component of the RNA induced silencing complex. Hc-Pro has been shown to interact with AGO1 (Ivanov et al, 2016), which, as discussed above, is recruited to the CBC in conditions of TOR inhibition, that are known to strongly repress WMV potyvirus accumulation (Ouibrahim et al, 2015). Overall, this suggest that AGO1, guided by antiviral siRNA, is involved in translational repression of the viral genomic RNA and that Hc-Pro may counteract this mechanism during normal infections. When TOR is inhibited the resulting over-recruitment of AGO1 to the CBC to drive general translational repression would also override the action of Hc-Pro, leading to virus elimination by RNA silencing. Interestingly, the Potato virus A HC-pro component, a viral suppressor of RNA silencing, induces the formation of RNA granules containing the ribosomal protein P0 (also identified as a TOR target, Dobrenel et al. 2016), AGO1, VCS and eIF(iso)4E, which are involved in the stimulation of PVA translation (Hafrén et al. 2015). This suggests that a large protein complex containing CBC-interacting components like eIF(iso)4E, AGO1 and VCS could be formed upon TOR inhibition and/or viral infection.

## Conclusions

In this work we analyze the data from 11 deep proteomic analysis of Arabidopsis CBC performed under different conditions. Most proteins were detected only in single treatment and with minimal peptides representation, raising caution about their physiological relevance in association with the CBC. However, a subset of proteins are reproducibly and highly represented in several experiments, and among them known components of the CBC such as translation initiation factors of the eIF4E, eIF4G and eIF3 families, validating the global approach. We also confirm the association of several proteins to the CBC that were identified by Bush and Doonan (2009) or subsequently (Patrick et al, 2018; Toribio et al. 2019). This wide analysis also highlights new potential links between the CBC and mRNA metabolic pathways such as RNA silencing and the TOR pathway that could be tested experimentally. In contrast, the LARP1 protein, an established translational regulator presumed to be associated to the cap (Scarpin et al, 2020), is only found in the subset of experiments L3 and L4, which represent the deepest analysis in our set. This may indicate that if LARP1 binds the CBC, which has not been demonstrated in plants, then its affinity might be too weak to for retention during the purification procedure. A new finding is the association of AGO1 to the plant CBC when TOR activity is repressed. Following TOR inhibition, AGO1 may adhere more voraciously to the mRNA cap or to eIF4E, orientating RNA silencing from specific RNA repression in association with diverse small RNA guides towards global translational repression. From a TOR centered view, this would be an energetically favorable situation. Whatever the exact mechanism that links TOR dependent translational repression and RNA silencing, it is remarkable that another component of RNA silencing, the AGO1 associated SGS3 double stranded RNA binding protein also interacts with the TOR complex (van Leene et al 2019).

The activity of many proteins acting at the CBC, like in all dynamic macromolecular complexes, are likely to be modulated by post-translational modifications. Indeed, several plant translation initiation factors are known to be phosphorylated in response to changing physiological conditions (Browning and Bailey-Serres, 2015). We anticipate that the data made available in this work will facilitate the systematic study of the central role of translational regulations in plants.

## Acknowledgements

This work was partly funded by the ANR program TranslaTOR (ANR-11-BSV6-0010) to AM, CM and CR and LabEx Saclay Plant Sciences-SPS (ANR-10-LABX-0040-SPS) to AM and CM. We thank the proteomic platforms Edyp-Grenoble, PAPPSO-Moulon and IMM-Marseille for their help in experiment design and data analysis and Dr. Benjamin Field for corrections on the manuscript. The proteomic experiments performed in Grenoble were partially supported by Agence Nationale de la Recherche under projects ProFI (Proteomics French Infrastructure, ANR-10-INBS-08) and GRAL, a program from the Chemistry Biology Health (CBH) Graduate School of University Grenoble Alpes (ANR-17-EURE-0003).

## Authors contribution

AM, LO, CR, RL, MA, YC, MB, CL performed experiments, AM, LO, CL, CC, CM, CR designed experiments and analyzed the data, CM and CR wrote the manuscript.

## Supplemental data legends

**Supplemental data table 1:** Lists of the proteins which were identified in all the proteomic analyses of purified CBC. Experiments names correspond to the ones defined in Table I.

**Supplemental data tables 2-5** : Proteomic data of CBCs purified from Col0 (Table 2), eIF4E mutant (Table 3), TOR constitutive RNAi 35-7 line (Table 4), and Sepharose 4B control (Table 5, proteins identified from two gel slices for each conditions).

**Supplemental data tables 6-7** : Proteomic data of CBCs purified from Col0 (Table 6), and Col0 infected with Watermelon Mosaic Viruses (Table 7). Data from Sepharose4B binding proteins was extracted from Supplemental table 5.

**Supplemental data table 8**: Proteomic data of CBCs purified from ethanol induced alcA::Gus control, alcA::TOR RNAi 5.3, alcA::TOR RNAi6.3 lines.

**Supplemental data tables 9-10**: Proteomic data of CBCs purified from Col0 seedlings treated with the TOR inhibitor AZD-8055 (Table 9, L3) and from control Col0 treated with DMSO for 15 min (Table 10, L4).

## Notes

### Competing Interest Statement

The authors have declared no competing interest.

